# IL-27R signaling serves as immunological checkpoint for NK cells to promote hepatocellular carcinoma

**DOI:** 10.1101/2020.12.28.424635

**Authors:** Turan Aghayev, Iuliia O. Peshkova, Aleksandra M. Mazitova, Elizaveta K. Titerina, Aliia R. Fatkhullina, Kerry S. Campbell, Sergei I. Grivennikov, Ekaterina K. Koltsova

## Abstract

Hepatocellular carcinoma (HCC) is the most common form of liver cancer with poor survival and limited therapeutic options. HCC has different etiologies, typically associated with viral or carcinogenic insults or fatty liver disease and underlying chronic inflammation presents as a major unifying mechanism for tumor promotion. On the other hand, mechanisms of how inflammatory response can regulate anti-cancer immunity in HCC remain incompletely understood.

Interleukin (IL)-27 receptor signaling plays an anti-inflammatory role in a variety of infectious and chronic inflammatory diseases. Here, using genetic and pharmacological approaches we found that IL-27 receptor (IL-27R) signaling promotes HCC development *in vivo*. Genetic loss of IL-27R suppressed HCC development in both toxin/carcinogen-induced diethylnitrosamine (DEN) and non-alcoholic steatohepatitis (NASH)-driven models. Elevated expression of IL-27RA rendered poor prognosis to HCC patients. Mechanistically, the pro-tumorigenic effect was mediated by immunoregulatory role of IL-27R signaling within the tumor microenvironment, particularly the suppression of Natural killer (NK) cells. IL-27R ablation enhanced the accumulation and activation of cytotoxic NK cells during acute liver injury and in HCC tumors, while depletion of NK cells abrogated the effect of genetic IL-27R disruption.

Taken together, our data suggest an unexpected role of IL-27R signaling as a novel immunological checkpoint regulating NK cell activity and promoting development of HCC of different etiologies.

## Introduction

Hepatocellular carcinoma (HCC) is the most common form of liver cancer with poor survival rate and limited treatment options (1,2). While innovative therapeutic strategies targeting tumor microenvironment and immune contexture provide hope to many cancer patients (3,4), liver cancers remain poorly responsive to immunotherapy and continue to be highly lethal disease (2,5). For example, despite wide clinical use of T cell activation-based immunotherapies in several cancer types (6), only recently checkpoint inhibitors demonstrated potentially promising results in HCC (7). Preventive approaches aimed at hepatitis B vaccination and hepatitis C eradication could potentially curb new HCC cases, however liver cancers caused by environmental toxins and fatty liver disease are clearly on the rise (8,9).

Chronic liver inflammation induced by infections, alcohol or obesity drives chronic injury and promotes compensatory proliferation of transformed hepatocytes leading to enhanced HCC onset (10). While anti-cancer immune responses play an important tumor-restrictive role, it remains to be determined how they are regulated by the inflammatory entities in HCC (11).

Irrespective of its etiology, HCC development is accompanied by the accumulation of various immune cells, which could contribute to cancer progression via the production of pro-inflammatory cytokines such as IL-6, IL-1, TNF and IL-17 (12). Cytokine signaling can enable HCC growth by activating proto-oncogenic transcription factors such as NF-κB and STAT3 in transformed hepatocytes and HCC cells (10,12–14). On the other hand, the presence of cytotoxic or IFN-γ producing T cells could suppress tumor growth (15). Moreover, liver microenvironment is uniquely enriched in Natural Killer (NK) cells, capable of tissue immunosurveillance and anti-tumorigenic functions potentially impacting liver cancer development (16,17). During HCC development number and activation of liver NK cells gradually declines due to yet unidentified mechanism probably related to specific signals originating from tumor microenvironment and chronic exposure to cancer cells (18). The mechanisms that drive functional suppression of NK cells and undermine such innate anti-cancer immune responses remains incompletely understood.

Interleukin (IL)-27 is a member of the IL-6/IL-12 cytokine superfamily and is an important regulator of immune responses (19). The IL-27 receptor (IL-27R) is expressed by some non-hematopoietic cells and by multiple immune cell subsets, including NK cells (20). IL-27R signals primarily via STAT1 and STAT3 (19), amongst which STAT3 is hepatocyte-intrinsic driver of HCC (21). IL-27 was shown to have a broad anti-inflammatory role in infectious and chronic immune-mediated diseases (22–24). IL-27R ablation leads to elevated production of pro-inflammatory cytokines, including IL-17A and IL-6 (23,25), indicating that in inflammation-driven cancer, such as HCC, IL-27 cytokine could potentially play a tumor-restrictive, anti-inflammatory role. *In vitro* and *in vivo* experiments revealed that IL-27 can directly control LAG-3, TIM-3, PD-1 and TIGIT inhibitory molecules expression in T cells (26). While immunomodulatory effects by IL-27 were shown in various pathophysiological models (27–30), the context-dependent and cell type specific underlying mechanisms remain incompletely understood. The protective role of IL-27 and its receptor signaling in cancer has been suggested based on its anti-inflammatory role in inflammatory diseases and studies using subcutaneous cell line transplants (31–33), however the role of IL-27R signaling in cancer development *in vivo* has not been thoroughly addressed.

Here we found that genetic loss of IL-27R surprisingly suppressed liver cancer development in two different *in vivo* models of HCC, including diethylnitrosamine (DEN) carcinogen-driven and non-alcoholic steatohepatitis (NASH)-driven HCC. Analysis of human HCC revealed that patients with high IL-27R expression display poor survival and have more advanced stages of the HCC. Mechanistically, we found that IL-27R signaling limited the accumulation and activation of natural killer (NK) cells in HCC tumors and IL-27R genetic ablation relieved the inhibition and resulted in infiltration of activated NK cells and reduction in HCC burden. IL-27R signaling repressed expression of NK–activating stress ligands on tumor cells; expression of NK-cell recruiting chemokines and NK cell activating receptors. Inactivation of IL-27R signaling in turn relieved that suppressive effect and enhanced NK cells anti-tumor immune responses *in vivo*. The dependence of IL-27R signaling driven immunoregulatory effects in HCC on NK cells was further established in experiments, where NK cell depletion reverted the effect of IL-27R inactivation on HCC development. Taken together, our data suggest that inactivation of IL-27R signaling may enhance NK cell accumulation and cytotoxicity and reverse their exhaustion in HCC, providing new therapeutic opportunities as well as preventive approaches in high risk patients with NASH and liver fibrosis set to progress to HCC.

## Results

### IL-27R signaling promotes HCC development in DEN-induced model

While the role of IL-27R signaling has been investigated in infectious and inflammatory diseases (34–38), little is known about its possible contribution to tumor development and progression *in vivo*. We first analyzed the expression of this cytokine receptor in human HCC tumors. IHC staining of human liver tissue with developed HCC revealed that IL-27R is expressed by infiltrating immune cells as well as hepatocytes (Fig.1A). Next, we evaluated the impact of *IL27RA* expression in human HCC tumors on disease-free survival using TCGA provisional data subjected to Kaplan-Meier analysis. We found that patients with high *IL27RA* expression exhibited poor disease-free survival and presented with more advanced stages of HCC, suggesting a potential cancer-promoting role of IL-27R signaling in HCC (Fig.1B, C). Such a role is nevertheless surprising as IL-27R signaling was previously shown to reduce the expression of IL-6, IL-17A and other inflammatory cytokines deemed pro-tumorigenic in HCC (12,23).

**Figure 1.**
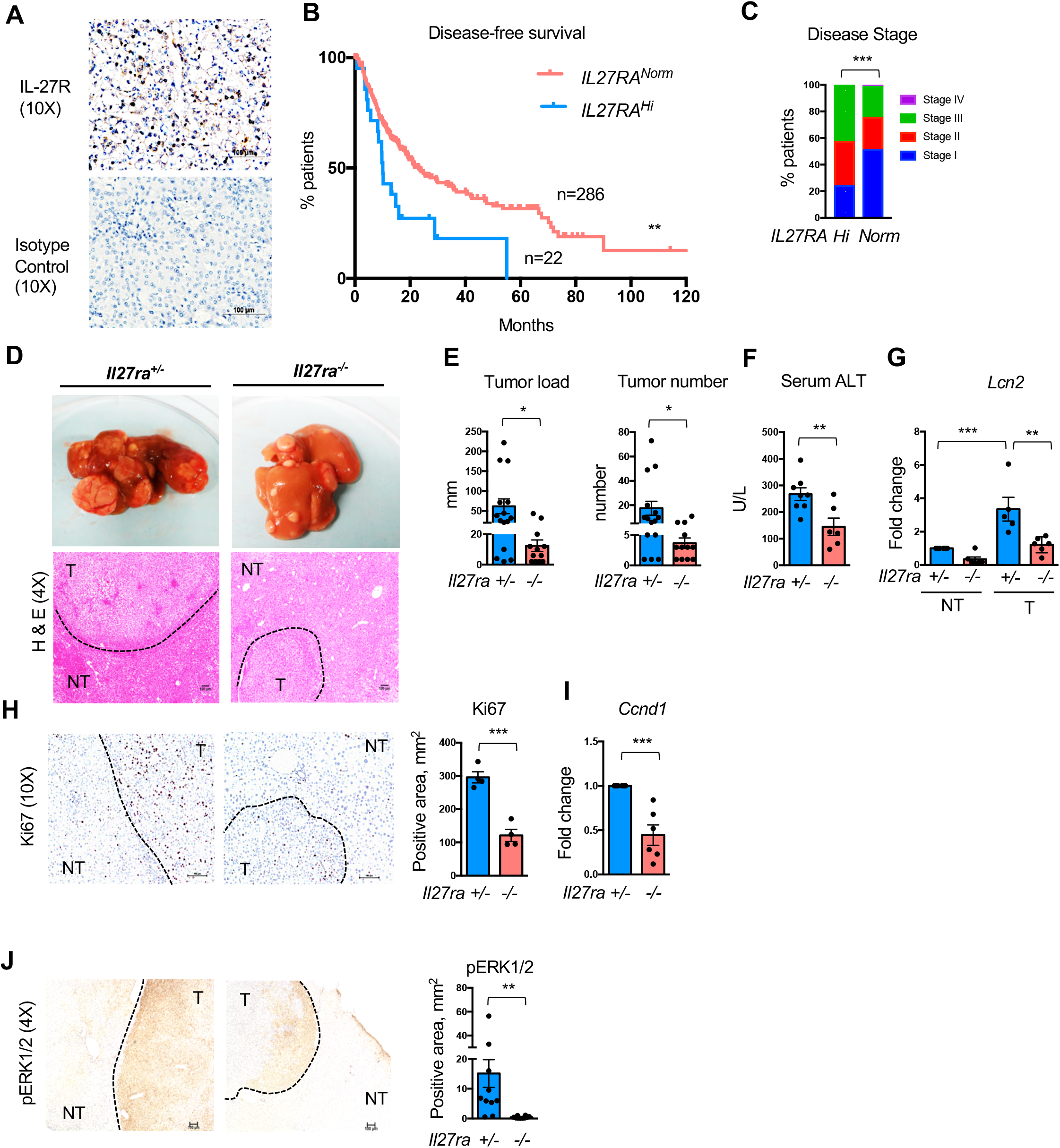
IL-27R signaling is implicated in liver cancer development. **(A)** Representative images of IL-27R expression in human HCC as determined by IHC staining. **(B)** TCGA provisional data on disease-free Kaplan-Meier estimate (disease free status since initial treatment) relative to *IL27RA* expression. **(C)** Correlation between *IL27RA* expression and HCC stage in patients based on TCGA data. **(D)** *Il27ra*^+/−^ and *Il27ra*^−/−^ mice received DEN at postnatal day 15. Cancer development was analyzed at 10 months of age. Representative images of macroscopic and microscopic view of livers with developed tumors. **(E)** Tumor load and tumor number in *Il27ra*^+/−^ (n=15) and *Il27ra*^−/−^ (n=12) mice. **(F)** Concentration of ALT in sera from*Il27ra*^+/−^ (n=8) and *Il27ra*^−/−^ (n=6) tumor-bearing 10-month-old mice. (G) Relative gene expression of inflammatory marker *Lcn2* in non-tumor and tumor tissues from *Il27ra*^+/−^ (n=6) and *Il27ra*^−/−^ (n=6) mice. Gene expression was first normalized to *Rpl32* then to gene expression in non-tumor tissue from *Il27ra*^+/−^ mice. **(H)** Representative images and quantification of Ki67 staining of HCC sections from DEN-treated *Il27ra*^+/−^ (n=4) and *Il27ra*^−/−^ (n=4) mice. **(I)** Relative gene expression of proliferation marker Cyclin D (*Ccnd1)* in tumors from *Il27ra*^+/−^ (n=6) and *Il27ra*^−/−^ (n=6) mice. Gene expression was first normalized to *Rpl32* then to gene expression in tumors from *Il27ra*^+/−^ mice. **(J)** Representative images and quantification of pERK1/2 staining of HCC sections from DEN-received *Il27ra*^+/−^ (n=4) and *Il27ra*^−/−^ (n=4) mice. **p<0.01, log-rank (Mantel-Cox) test (B); ***p<0.001, Chi-square test (C). Data are mean ± SEM from at least 3 independent experiments. *p<0.05, **p<0.01, ***p<0.001, unpaired Student’s t-test (two-tailed) (E, F, H-J); **p<0.01, ***p<0.001, Tukey’s multiple comparisons test (G).

To directly evaluate the potential role of IL-27R signaling in HCC we used a genetic approach in a well-established mouse model of HCC driven by the toxin/carcinogen DEN (10,39,40). To exclude any confounding influence of genetics or microbiota, we used cage-mate and littermate controls. Male *Il27ra*^+/−^ or *Il27ra*^−/−^ mice were injected with 25 mg/kg of DEN i.p. at day 15 after birth and tumor development was assessed at 10 months of age. Given previous observations that IL-27 dampens inflammatory signaling (20), we expected that IL-27R ablation would exacerbate inflammation and promote *in vivo* HCC development. Unexpectedly, we found that HCC tumorigenesis was markedly reduced in *Il27ra*^−/−^ mice compared to *Il27ra*^+/−^ controls (Fig.1D, E). Body weight of tumor-bearing mice was not affected by IL-27R deficiency (Suppl. Fig.1A). Serum level of ALT, a marker of liver damage reflecting HCC formation, was significantly lower in *Il27ra*^−/−^ mice compared to their *Il27ra*^+/−^ tumor bearing counterparts suggesting preserved liver function in the absence of IL-27R (Fig.1F). No difference was detected in serum Albumin, Globulin or total bilirubin (Suppl. Fig.1B). In line with limited liver damage, HCC tumors from IL-27R deficient mice were also characterized by lower expression of the inflammatory marker *Lcn2* (Fig.1G). Tumors of IL-27R deficient mice were characterized by reduced proliferation as determined by Ki67 staining (Fig.1H) and cyclin D1 (*Ccnd1)* expression (Fig.1I). Consistently with lower proliferation and inflammation/injury, activation of kinase ERK1/2 was also reduced in IL-27R deficient tumors (Fig.1J). To determine if limited proliferation is a broader representation of IL-27R-dependent liver regeneration capacity, we compared liver regeneration in IL-27R deficient and sufficient mice. Partial hepatectomy of 2/3 of liver tissue revealed no significant differences between *Il27ra*^−/−^ or control *Il27ra*^+/−^ mice in the ability to regenerate liver mass, suggesting that IL-27R regulates HCC development independent of its effects on liver regeneration (Suppl. Fig.2A). *In vitro* stimulation of DEN-derived primary HCC cells with recombinant IL-27 (rIL-27) was also unable to change HCC cell growth in an *in vitro* clonogenic assay (Suppl. Fig.2B), despite induction of STAT3 phosphorylation in HCC cells (Suppl. Fig.2C).

Collectively, our data demonstrate that IL-27R signaling promotes HCC in the carcinogen-induced model *in vivo*.

### IL-27R signaling restricts NK cell accumulation in HCC

HCC progression is regulated by various immune cells, accumulating in tumor and non-tumor tissue (12). In order to elucidate the potential mechanisms by which IL-27R signaling controls tumor growth we first conducted NanoString gene expression analysis on isolated tumors from IL-27R deficient and proficient animals. Among the most notable changes we found an increase in NK-specific markers and overall upregulation of the NK-mediated cytotoxicity pathway as defined by KEGG analysis in tumors of *Il27ra*^−/−^ mice, implying a potential role of IL-27R-dependent NK cell regulation in the control of HCC (Fig.2A, B).

**Figure 2.**
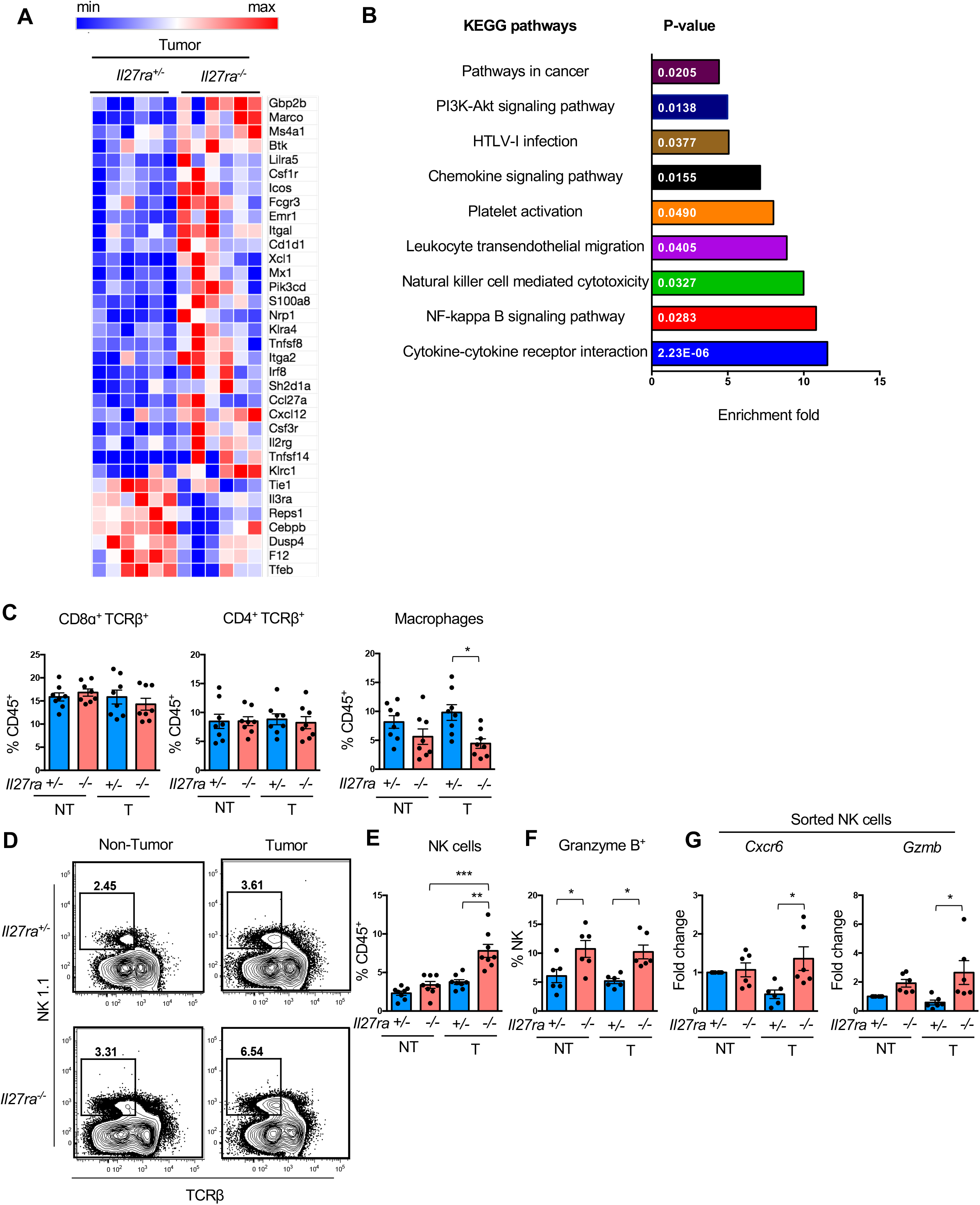
IL-27R signaling impairs NK cell accumulation and cytotoxicity. Heatmap (A) and KEGG pathway analysis (B) of differentially expressed genes in tumors from DEN-treated *Il27ra*^+/−^ (n=6) and *Il27ra*^−/−^ (n=6) mice as determined by NanoString (p<0.05). Single cell suspensions of non-tumor and tumor tissues from *Il27ra*^+/−^ (n=8) and *Il27ra*^−/−^ (n=8) DEN-injected 10-month-old mice were stained for Live/Dead, CD45, TCRβ, CD8α, CD4, NK1.1, CD11b, Ly6G, Ly6C, F4/80, Granzyme B and analyzed by FACS. **(C)** Percentage of CD8α^+^ TCRβ^+^ and CD4^+^ TCRβ^+^ cells, and macrophages. **(D)** Representative FACS plots of NK cells in non-tumor and tumor tissue. **(E)** Percentage of NK cells and (F) Granzyme B+ NK cells as determined by intracellular staining of non-tumor and tumor tissue from *Il27ra*^+/−^ (n=6) and *Il27ra*^−/−^ (n=6) mice. (G) Relative gene expression of *Cxcr6* and *Gzmb* in NK cells sorted by FACS from non-tumor and tumor tissue from *Il27ra*^+/−^ (n=6) and *Il27ra*^−/−^ (n=6) mice. Gene expression was first normalized to *Rpl32* then to gene expression of NK cells in non-tumor tissue from *Il27ra*^+/−^ mice. Data are mean ± SEM from at least 3 independent experiments. *p<0.05. **p<0.01, ***p<0.001, Tukey’s multiple comparisons test.

The analysis of immune cell composition in tumor and adjacent non-tumor tissue by flow cytometry revealed no differences in CD4 or CD8 T cell infiltration with some reduction of macrophages in tumors of *Il27ra*^−/−^ mice (Fig.2C). However, in concert with our gene expression data, we found a significant increase in NK cells in tumors of IL-27R deficient mice compared to IL-27R sufficient controls (Fig.2D, E). These NK cells were characterized by elevated Granzyme B production as determined by flow cytometry and Q-RT-PCR analysis on FACS sorted NK cells from HCC tumors and adjacent normal tissue (Fig.2F, G), suggesting their enhanced activation and cytolytic potential. Sorted NK cells also expressed higher level of *Cxcr6, a* chemokine receptor that is characteristically expressed on hepatic NK cells and regulates their accumulation (41,42) (Fig.2G).

### IL-27R acts as immunological checkpoint taming NK cell activation in HCC

NK cell activation is controlled by multiple activating and inhibitory receptors, which regulate their “active” versus “inactive” state (43). NKG2D activating receptor engagement by “stress molecule” ligands on target cells increases NK cell activity (44–46), while inhibitory receptor Ly-49C engages MHC class I (H-2K^b^ and H-2D^b^) on target cells and dampens NK cell activation (45,47).

Therefore, next we analyzed the expression of NK cell activating and inhibitory receptors in IL-27R sufficient and deficient mice. We found upregulation of activating receptor NKG2D (Fig.3A, B) and NKG2A, a marker of less mature NK cells that educates CXCR6^+^ liver-resident NK cells (41,48) (Fig.3C) and downregulation of inhibitory receptor Ly-49C on NK cells (49) isolated from HCC tumors of *Il27ra*^−/−^ mice (Fig.3D, E).

**Figure 3.**
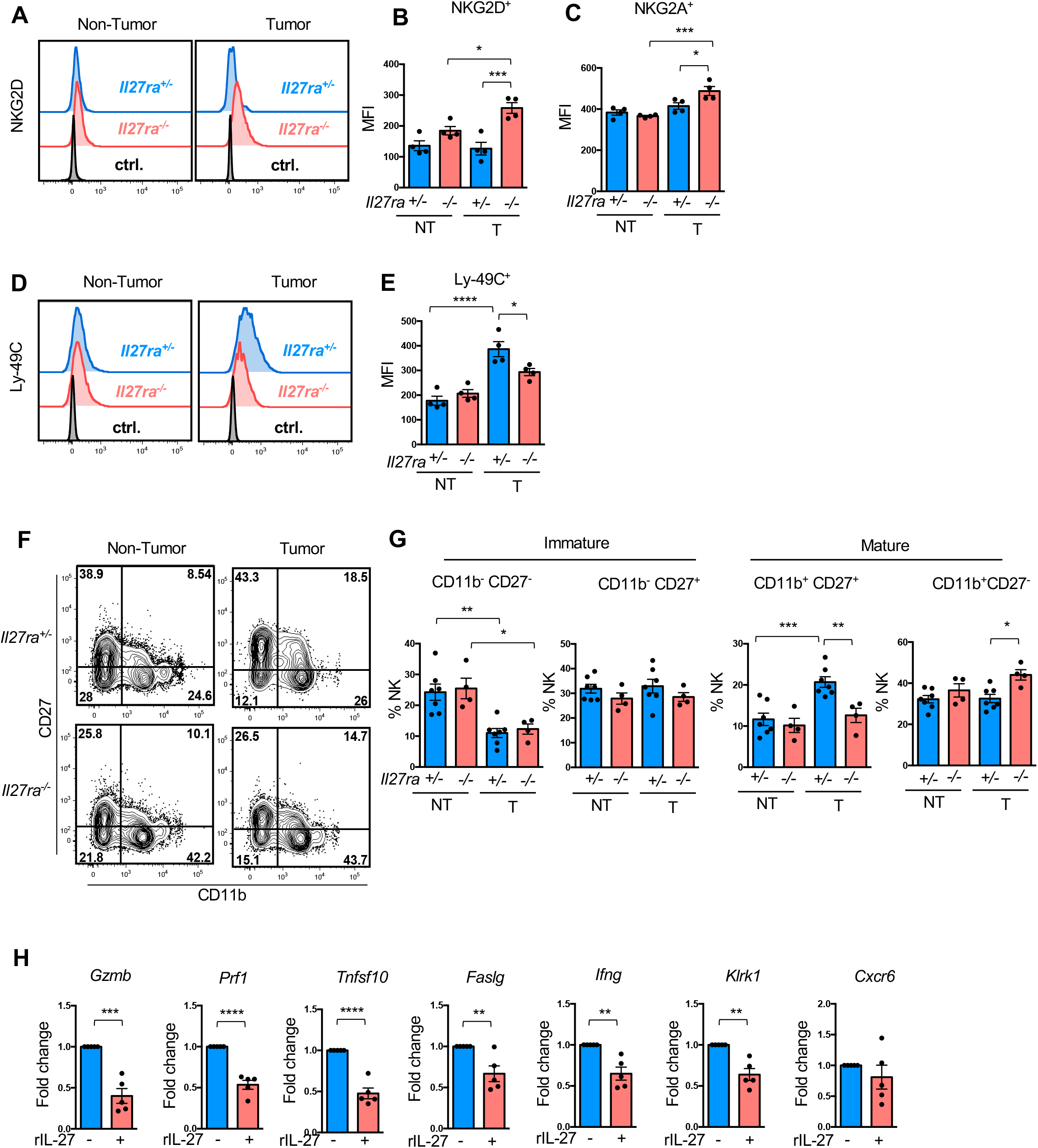
IL-27R signaling augments NK cell activation and maturation. Single cell suspensions of non-tumor and tumor tissue from DEN-injected *Il27ra*^+/−^ (n=4) and *Il27ra*^−/−^ (n=4) 10-month-old mice were stained for Live/Dead, CD45, TCRβ, NK1.1, CD49a, CD49b, NKG2D and NKG2A. **(A)** Representative histogram of NKG2D activating receptor expression. (B) Mean fluorescence intensity and percentage of NKG2D+CD49b+ NK cells in non-tumor and tumor tissue. (C) Mean fluorescence intensity and percentage of NKG2A^+^CD49b^+^ NK cells in non-tumor and tumor tissue. (D) Representative histogram of Ly-49C inhibitory receptor expression. (E) Mean fluorescence intensity and percentage of Ly-49C^+^CD49b^+^ NK cells in non-tumor and tumor tissue. Single cell suspensions of non-tumor and tumor tissue from DEN-injected *Il27ra*^+/−^ (n=7) and *Il27ra*^−/−^ (n=4) 10-month-old mice were stained for Live/Dead, CD45, TCRβ, NK1.1, CD11b, CD27 and analyzed by FACS. **(F)** Representative FACS plots of NK-cell maturation based on CD11b and CD27 expression. **(G)** Percentage of immature (CD11b^−^CD27^−^ and CD11b^−^CD27^+^) and mature (cytotoxic/cytokine secreting CD11b^+^CD27^+^ and terminally mature CD11b^+^CD27^−^) NK cells. (H) Relative gene expression of *Gzmb*, *Prf1*, *Ifng*, *Klrk1*, *Tnfsf10*, *Faslg* and *Cxcr6* in NK cells purified by MACS from spleens of WT mice (n=5) and stimulated *in vitro* with rIL-27. Gene expression was first normalized to *Klrb1c* then to gene expression in untreated condition. Data are mean ± SEM from at least 3 independent experiments. *p<0.05. **p<0.01, ***p<0.001, Tukey’s multiple comparisons test (B, C, E, G); **p<0.01, ***p<0.001, ****p<0.0001, unpaired Student’s t-test (two-tailed) (H).

Next, we analyzed whether IL-27R signaling affects the maturation of NK cells. We found elevated presence of terminally differentiated mature CD11b^+^CD27^−^ NK cells (50) in HCC tumors from *Il27ra*^−/−^ mice, suggesting potential inhibiting role of IL-27R on NK cell maturation (Fig.3F, G). In contrast, more of cytokine-producing mature CD11b^+^CD27^+^ NK cells (50) were found in tumors of *Il27ra*^+/−^ control animals (Fig.3G). No significant changes in NKG2D, NKG2A or Ly-49C expression were detected in blood or spleen suggesting their site-specific activation (Suppl. Fig.3A,B)

To further understand functional implication of IL-27 signaling in controlling NK cell activity we performed an *in vivo* cytotoxicity assay using RMA-S (sensitive to NK killing) and RMA (insensitive) tumor cell lines. Cells were dye-labeled and injected i.p. at 1:1 ratio into naïve *Il27ra*^−/−^ and *Il27ra*^+/−^ mice. After 48h, flow cytometry analysis of the peritoneal lavage revealed a reduced percentage of RMA-S cells in *Il27ra*^−/−^ mice compared to *Il27ra*^+/−^ controls, suggesting enhanced NK cell cytotoxicity towards the sensitive line in the absence of IL-27R (Suppl. Fig.3C-E). Taken together, these data suggest that IL-27R signaling is implicated into the regulation of NK cell accumulation, activation and cytotoxicity during HCC development.

To test whether IL-27 can directly regulate NK cells, sorted NK cells from wild type mice were stimulated *in vitro* with rIL-27. We found that IL-27 was able to suppress the expression of various cytotoxic and activating molecules including granzyme B (*Gzmb*), perforin (*Prf1*), TRAIL (*Tnfsf10*), FasL (*Faslg*) as well as IFN-γ (*Ifng*) and NKG2D (*Klrk1*) (Fig.3H) without affecting cell survival, implying its direct role in suppressing NK effector phenotype.

In order to further characterize the link between liver injury, IL-27R signaling and NK cell accumulation, we analyzed livers from mice subjected to acute carcinogen-induced liver injury. Eight-week-old *Il27ra*^+/−^ and *Il27ra*^−/−^ mice were administered with 100 mg/kg DEN and gene expression in the liver was analyzed 48 hours later by Q-RT-PCR. We found strongly elevated expression *of Cxcl9* and *Cxcl10* chemokines as well as Prf1, *Gzmb*, and *Ifng* in the livers of DEN-treated *Il27ra*^−/−^ mice (Suppl. Fig.3F, G), indicating that IL-27R controls NK cell recruitment and/or activation during early and late stages of HCC development and may counteract early immunosurveillance exerted at the level of tumor seeds.

### IL-27R signaling regulates NK cell activation through repression of “stress ligand” expression

Activation of NK cells requires engagement of activating receptors on NK cells with corresponding stress-induced ligands on target cells. In particular, NK cell activation is dependent on RAE-1 and H60 families of activating ligands (51,52) and is further enhanced by MHC I downregulation, a “missing self” signal on target cells (53). We analyzed the expression of stress ligands, *Raet1a* and *H60b* in normal or tumor tissues of mice with DEN-induced HCC. Q-RT-PCR analysis revealed a significant upregulation of both *Raet1a* and *H60b* expression in tumors of *Il27ra*^−/−^ mice compared to *Il27ra*^+/−^ controls (Fig.4A). Conversely, rIL-27 was able to suppress *Raet1a* and *H60b* expression in the DEN-derived primary HCC cell line (Fig.4B). Furthermore, a significant downregulation of surface MHC I expression on tumor cells was observed in tumors from *Il27ra*^−/−^ mice compared to *Il27ra*^+/−^ controls (Fig.4C), along with a reduction in *Tap1* expression, a regulator of peptide loading on MHC class I molecules (Fig.4D). These data imply that IL-27R signaling promotes expression of MHC I and represses upregulation of NK–triggering ‘‘stress ligands”, thereby serving as a direct and indirect immunoregulator of NK cell activity and NK cell mediated anti-tumor responses.

**Figure 4.**
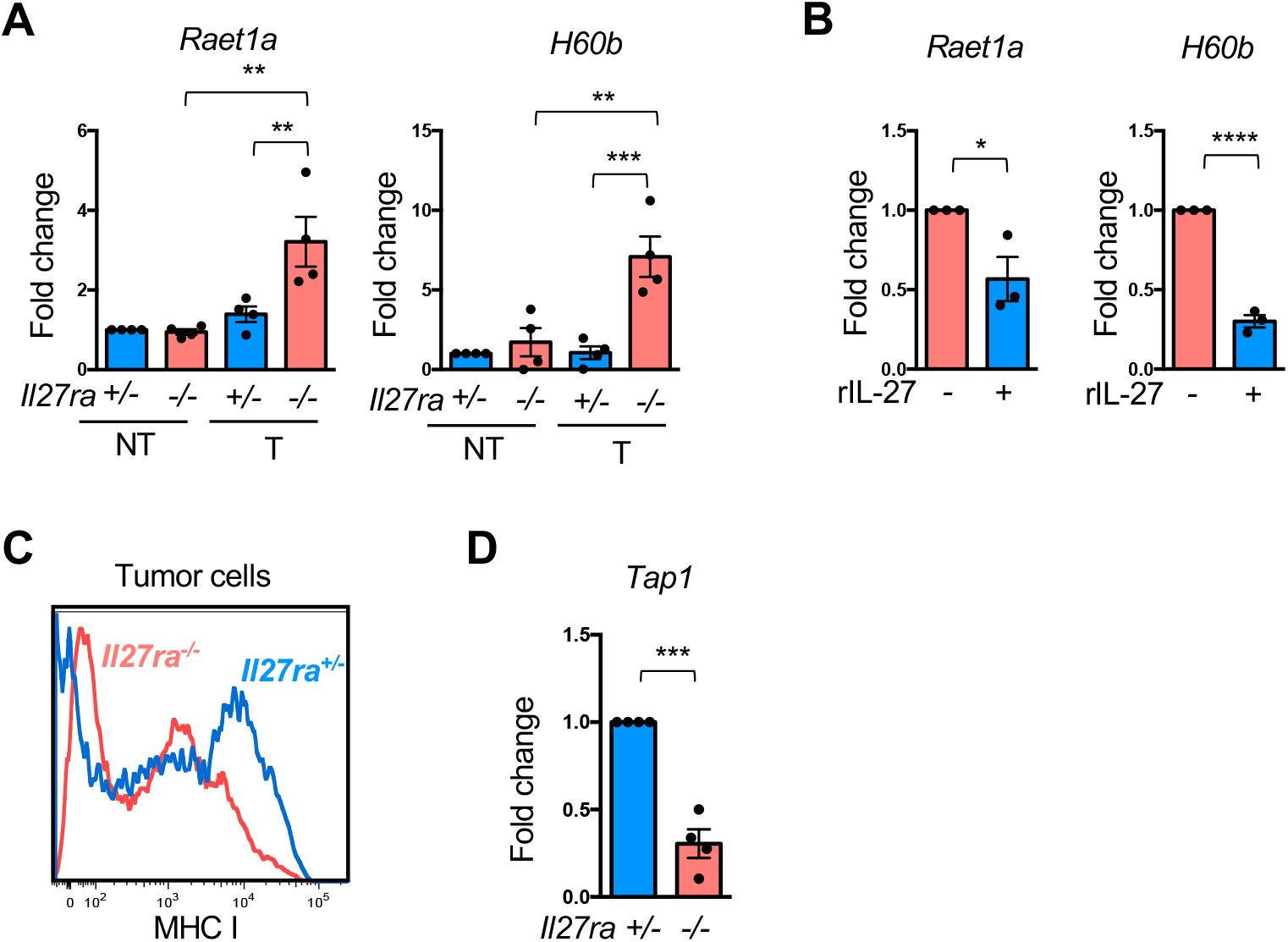
IL-27R signaling controls stress molecules and MHC I expression. **(A)** Relative gene expression of stress molecules *Raet1a* and *H60b* in non-tumor and tumor tissue from DEN-injected *Il27ra*^+/−^ (n=4) and *Il27ra*^−/−^ (n=4) mice. Gene expression was first normalized to *Rpl32* then to gene expression in non-tumor tissue from *Il27ra*^+/−^ mice. **(B)** Relative gene expression of *Raet1a* and *H60b* in DEN-derived HCC cells treated *in vitro* with rIL-27. Gene expression was first normalized to *Rpl32* then to gene expression in untreated condition. Single cell suspensions of tumor tissue from DEN-injected *Il27ra*^+/−^ and *Il27ra*^−/−^ 10-month-old mice were stained for Live/Dead, CD45, CD11b, CD31, TER-119, H-2K^b^ and analyzed by FACS. **(C)** Representative histogram of MHC I expression on tumor cells from *Il27ra*^+/−^ and *Il27ra*^−/−^ mice. **(D)** Relative gene expression of MHC I processing gene *Tap1* in tumor tissue of *Il27ra*^+/−^ (n=4) and *Il27ra*^−/−^ (n=4) mice. Gene expression was first normalized to *Rpl32* then to that in tumor tissue of *Il27ra*^+/−^ mice. Data are mean ± SEM from at least 3 independent experiments. Tukey’s multiple comparisons test (A); *p<0.05. **p<0.01, ***p<0.001., unpaired Student’s t-test (two-tailed) (B, D).

### IL-27R signaling drives NASH-induced HCC

While the incidence of liver cancer caused by Hepatitis B and C declines due to efficient therapies and vaccines (54), obesity-driven HCC is on the rise (8,9,55). Obesity drives the development of non-alcoholic steatohepatitis (NASH), strongly promoting HCC (56). To complement our studies in toxin/mutagen-induced DEN model, we next sought to generalize our observations using a NASH-dependent model of HCC. We crossed *Il27ra*^−/−^ mice *to MUP-uPA* mice where the *uPA* (urokinase plasminogen activator) transgene is controlled by mature hepatocyte-specific promoter (*MUP*). *MUP-uPA* expression combined with Western diet (WD) feeding drives strong fibrosis and spontaneous HCC development and overall faithfully models NASH-driven HCC in humans (57–59). *MUP-uPA*^+^*Il27ra*^−/−^ and *MUP-uPA*^+^*Il27ra*^+/−^ mice were fed the WD starting at 8 weeks of age for a total of 8 months. We did not find any significant difference in body weight (Suppl. Fig.4). Similar to observations in the DEN-induced HCC model, IL-27R deficient *MUP-uPA*^+^ mice were largely protected from HCC development (Fig.5A, B). Tumors from *MUP-uPA*^+^*Il27ra*^−/−^ mice were characterized by the reduction of *Ccnd1* expression, implying reduced proliferative capacity of tumors in the absence of IL-27R signaling (Fig.5C). We also observed a reduction of collagen content in livers of *MUP-uPA*^+^*Il27ra*^−/−^ mice, suggesting limited fibrosis in the absence of IL-27R signaling (Fig. 5D). The level of *Lcn2* was decreased in tumors of IL-27R deficient *MUP-uPA*^+^ mice (Fig.5E).

**Figure 5.**
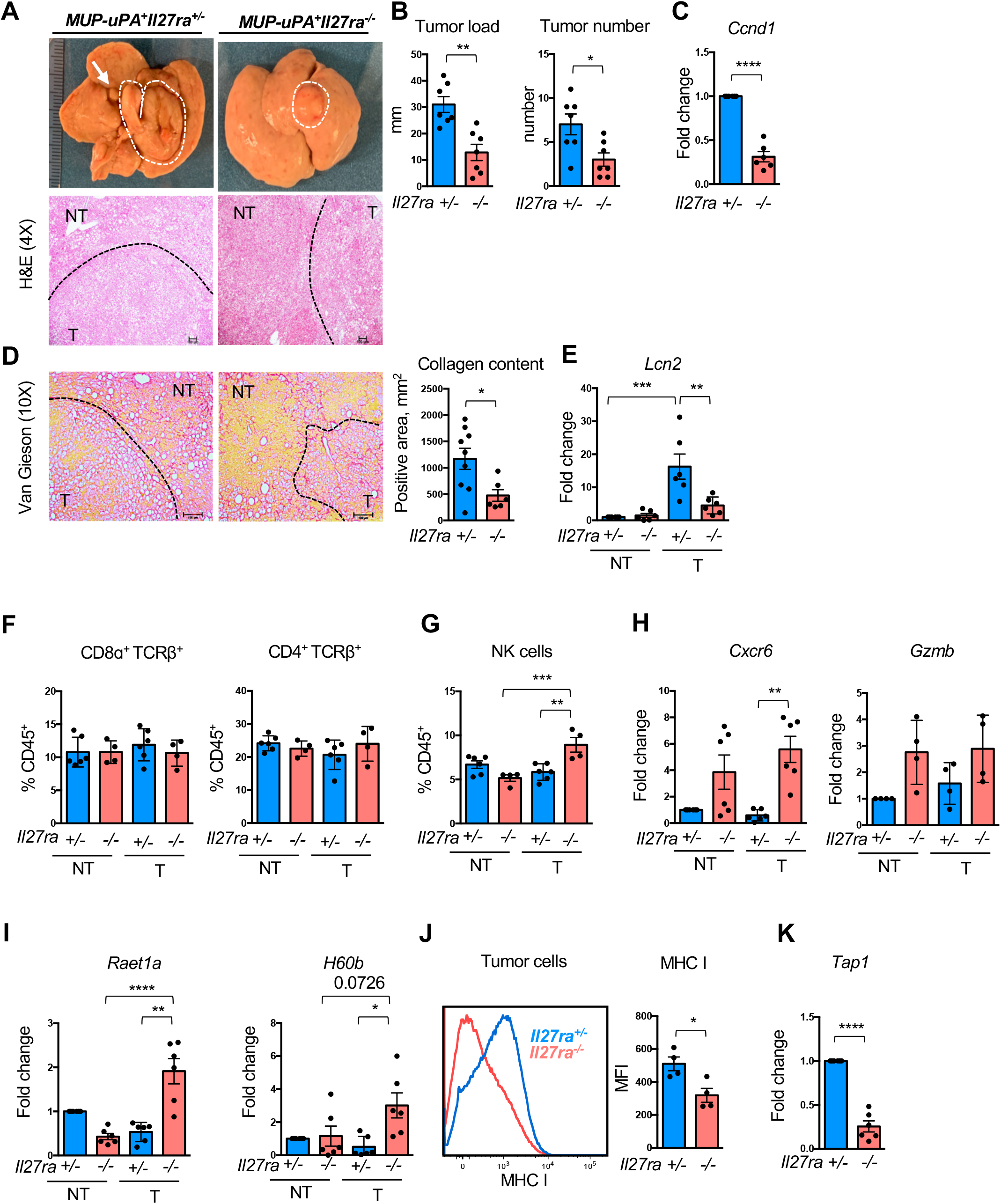
IL-27R signaling suppresses HCC tumor growth in NASH model. 8-week-old *MUP-uPA*^+^*Il27ra*^+/−^ and *MUP-uPA*^+^*Il27ra*^−/−^ mice were fed a WD for 8 months. **(A)** Representative images of macroscopic and microscopic view of livers with developed tumors. **(B)** Tumor load and tumor number of *MUP-uPA*^+^*Il27ra*^+/−^ (n=7) and *MUP-uPA*^+^*Il27ra*^−/−^ (n=7) male mice. (C) Relative gene expression of *Ccnd1* in tumor tissue from *MUP-uPA*^+^ *l27ra*^+/−^ (n=6) and *MUP-uPA*^+^*Il27ra*^−/−^ (n=6) male mice. Gene expression was first normalized to *Rpl32* then to gene expression in tumors from *MUP-uPA*^+^*Il27ra*^+/−^ mice. **(D)** Representative images and quantification of collagen content (collagen-red, other tissues-yellow) determined by Van Gieson staining of liver sections from *MUP-uPA*^+^*Il27ra*^+/−^ (n=3) and *MUP-uPA*^+^*Il27ra*^−/−^ (n=3) male mice. (E) Relative gene expression of *Lcn2* in non-tumor and tumor tissue from *MUP-uPA*^+^*Il27ra*^+/−^ (n=6) and *MUP-uPA*^+^*Il27ra*^−/−^ (n=6) female and male mice. Gene expression was first normalized to *Rpl32* then to gene expression in non-tumor tissue from *MUP-uPA*^+^*Il27ra*^+/−^ mice. Single cell suspensions of non-tumor and tumor tissue of *MUP-uPA*^+^*Il27ra*^+/−^ (n=6) and *MUP-uPA*^+^*Il27ra*^−/−^ (n=4) WD-fed 10-month-old female and male mice were stained for Live/Dead, CD45, TCRβ, CD8α, CD4, NK1.1 and analyzed by FACS. **(F)** Percentage of CD8α^+^TCRβ^+^ and CD4^+^TCRβ^+^ cells. **(G)** Percentage of NK cells. Relative gene expression of *Cxcr6*, *Gzmb* (H) and *Raet1a*, *H60b* (I) in non-tumor and tumor tissue from *MUP-uPA*^+^*Il27ra*^+/−^ (n=6) and *MUP-uPA*^+^*Il27ra*^−/−^ (n=6) female and male mice. Gene expression was first normalized to *Rpl32* then to gene expression in non-tumor tissue from *MUP-uPA*^+^*Il27ra*^+/−^ mice. Single cell suspensions of tumor tissue from WD-fed *MUP-uPA*^+^*Il27ra*^+/−^ (n=4) and *MUP-uPA*^+^*Il27ra*^−/−^ (n=4) 10-month-old female and male mice were stained for Live/Dead, CD45, CD11b, CD31, TER-119, MHC I and analyzed by FACS. **(J)** Representative histogram and mean fluorescent intensity of MHC I expression on tumor cells. **(K)** Relative gene expression of *Tap1* in tumors from *MUP-uPA*^+^*Il27ra*^+/−^ (n=6) and *MUP-uPA*^+^*Il27ra*^−/−^ (n=6) female and male mice. Gene expression was first normalized to *Rpl32* then to that in tumor tissue from *MUP-uPA*^+^*Il27ra*^+/−^ mice. Data are mean ± SEM from at least 3 independent experiments. *p<0.05. **p<0.01, ***p<0.001, ****p<0.0001, unpaired Student’s t-test (two-tailed) (B-D, J, K); Tukey’s multiple comparisons test (E, G-I).

The analysis of immune cell composition revealed no changes in CD4 or CD8 T cells (Fig.5F), but elevated accumulation of NK cells in tumors of *MUP-uPA*^+^*Il27ra*^−/−^ mice (Fig.5G). CXCR6 (*Cxcr6*) and granzyme B (*Gzmb*) expression was also upregulated in the absence of IL-27R as determined by Q-RT-PCR (Fig.5H). Consistent with the results in DEN model, we also detected heightened expression of *Raet1a* and *H60b* in tumors of IL-27R deficient MUP-uPA^+^ mice (Fig.5I) while MHC I and *Tap1* were downregulated in tumors of mice lacking IL-27R (Fig.5J, K). These data further advanced a functional mechanistic link connecting IL-27R signaling to regulation of NK cell mediated anti-tumor immunity and HCC development in models of different etiology.

### Tumor promoting effect of IL-27R signaling is exerted through suppression of NK cell mediated anti-tumor immunity

Next, we sought to establish the “linearity” of the mechanistic link between IL-27R signaling, NK cell activity and HCC development. We first tested whether NK cell activity is essential for the anti-tumor effect of IL-27R ablation. We depleted NK cells using anti-NK1.1 antibody in DEN-treated *Il27ra*^−/−^ and *Il27ra*^+/−^ mice for total period of 5.5 months prior to tumor collection. Efficiency of NK depletion was confirmed by FACS analysis of blood, non-tumor and tumor tissues (Suppl. Fig.5). The depletion of NK cells enhanced tumor growth in *Il27ra*^−/−^ mice and eliminated the differences between *Il27ra*^−/−^ and *Il27ra*^+/−^ cohorts (Fig.6A, B), suggesting that NK cells is a key functional immune population regulated by IL-27R in the context of HCC tumor progression.

NKp46 is a natural cytotoxicity receptor expressed by NK cells (60). The analysis of NKp46 expression on liver NK cells revealed enrichment of NKp46+ NK cells in *Il27ra*^−/−^ mice compared to *Il27ra*^+/−^ controls (Fig. 6C). Moreover, *in vitro* stimulation of sorted NK cells with rIL-27 suppressed NKp46 (*Ncr1*) expression, suggesting that IL-27R signaling could directly regulate this cytotoxicity receptor expression and therefore impact functional activity of NK cells (Fig.6D). We therefore took an advantage of NKp46-GFP reporter strain (*Ncr1^gfp/gfp^*), where *Ncr1* gene is replaced by gene expressing green fluorescent protein (GFP), marking NK cells (61). Importantly, heterozygous or homozygous disruption of NKp46 in these mice affects expression levels of NKp46 and cell activation and differentiation (62). *Ncr1*^+/*gfp*^*Il27ra*^−/−^ and *Ncr1*^+/*gfp*^*Il27ra*^+/−^ mice were administered with DEN as previously described and tumor development was analyzed at 10 months of age. Similar to NK cell antibody depletion, reduced NK cell activation capacity in *Ncr1*^+/*gfp*^ mice ‘cancelled out’ anti-tumor effect of IL-27R deficiency and resulted in similar tumor load and number of tumors formed in IL-27R deficient and IL-27R sufficient mice (Fig.6E, F compare to Fig.1D, E). Importantly, heterozygous *Ncr1*^+/*gfp*^ status did not affect tumor development in IL-27R sufficient mice (Fig. 6E, F compare to Fig.1D, E). Therefore, the presence and the activation of NK cells is required for the anti-tumor effect of IL-27R deficiency and IL-27R signaling promotes HCC development by repressing NK cell activity.

**Figure 6.**
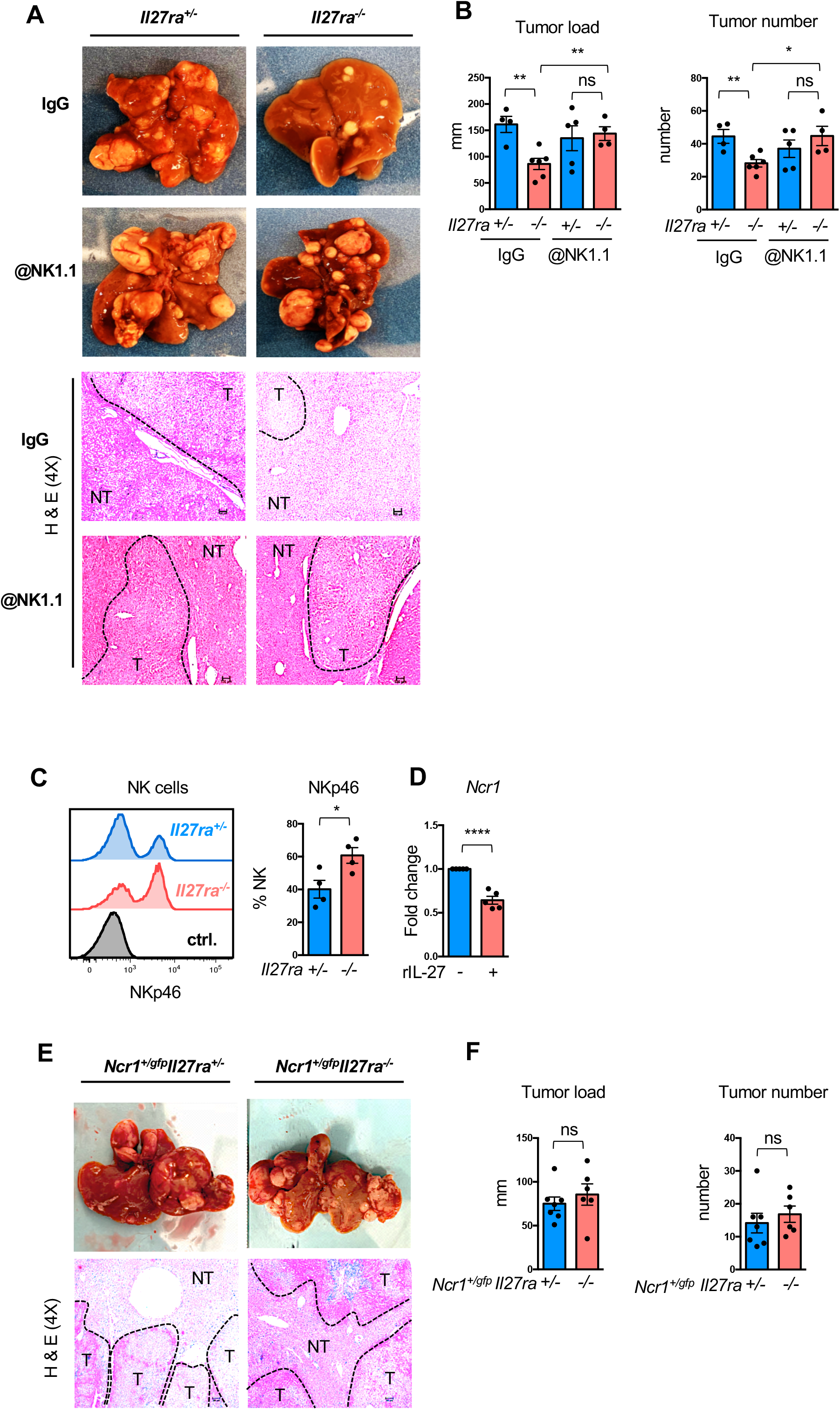
IL-27R signaling exerts its action via NK cells. DEN-treated *Il27ra*^+/−^ and *Il27ra*^−/−^ mice were administered with anti-NK1.1 and isotype control (IgG) for 5.5 months prior to tumor development analysis at 10 months. **(A)** Representative images of macroscopic and microscopic view of livers with developed tumors. (B) Tumor load and tumor number in IgG group: *Il27ra*^+/−^ (n=4), *Il27ra*^−/−^ (n=6) and @NK1.1 group: *Il27ra*^+/−^ (n=5), *Il27ra*^−/−^(n=4) mice. **(C)** Representative histogram and percentage of NKp46+ NK cells from livers of *Il27ra*^+/−^ (n=4) and *Il27ra*^−/−^ (n=4) mice as determined by FACS. (D) Relative gene expression of *Nrc1 (NKp46)* in NK cells purified by MACS from spleens of WT mice (n=5) and stimulated *in vitro* with rIL-27. Gene expression was first normalized to *Klrb1c* then to gene expression in untreated condition. **(E)** Representative images of macroscopic and microscopic view of livers with developed tumors from *Ncr1*^+/*gfp*^*Il27ra*^+/−^ and *Ncr1*^+/*gfp*^*Il27ra*^−/−^ DEN-treated mice analyzed at 10 months of age. **(F)** Tumor load and tumor number among *Ncr1*^+/*gfp*^*Il27ra*^+/−^ (n=7) and *Ncr1*^+/*gfp*^*Il27ra*^−/−^ (n=6) mice compared to *Il27ra*^+/−^ (n=15) and *Il27ra*^−/−^ (n=12) mice from the cohorts shown on Figure 1E. Data are mean ± SEM from at least 3 independent experiments. *p<0.05, **p<0.01, ****p<0.0001, unpaired Student’s t-test (two-tailed).

Overall, our data suggest that IL-27R signaling controls HCC tumor development *in vivo* as a new immunological checkpoint regulating NK cell activation and cytotoxicity. Particularly, we found that IL-27R signaling suppresses NK cell accumulation and activation by controlling the expression of activating and inhibitory receptors on NK cells, as well as MHC I and stress-induced ligands on cancer cells.

## Discussion

Primary liver cancer is the fifth most common cancer worldwide accounting for approximately 800,000 death each year (63). Despite efforts to curb HCC via implementation of vaccines against hepatitis B and drugs against hepatitis C, the incidence of liver cancer is sharply on the rise, because of the increased prevalence of fatty liver disease coupled with epidemic of obesity and type II diabetes, along with an increase in exposure to environmental toxins and pollutants. While chronic inflammation is known to contribute to liver cancer development, mechanisms of anti-cancer immune responses are not completely understood. Alike, as many patients with chronic liver disease are set to progress to HCC at some timepoint, approaches to curb HCC progression and new avenues for HCC therapies are in dire need.

IL-27 and its receptor signaling have been implicated in the regulation of inflammation in various acute and chronic inflammatory diseases (23,34,38,64,65). Meanwhile, the role of IL-27 in cancer development has not been extensively investigated, especially in faithful *in vivo* models. Various immune cells have been implicated in the control of tumor initiation and progression. The accumulation of polarized and inflammatory myeloid cells, Tregs, and the exclusion or exhaustion of conventional CD4 and CD8 T cells are associated with tumor progression (66). Similarly, chronic inflammation particularly manifested in elevated production of pro-tumorigenic cytokines and growth factors, is an underlying factor and a potent driver of cancer progression, including HCC (12). Hence, cytokines like IL-6, TNF or IL-17A were shown to be essential for HCC tumorigenesis in a variety of animal models (10,57), and elevated levels of IL-6 were identified as a key biomarker of liver disease-to-HCC progression (67). While IL-27 was originally described as a pro-inflammatory cytokine, subsequent studies demonstrated its largely anti-inflammatory role, placing IL-27 into the clan of immunoregulatory cytokines (68). We and others previously have shown that inactivation of IL-27 in chronic inflammatory diseases such as atherosclerosis results in enhanced inflammation and production of IL-6, IL-17A and other cytokines (23,24,37). Our initial hypotheses for this study therefore were that IL-27 would play an anti-inflammatory role in HCC, thereby reducing expression of key inflammatory tumor promoters, and that IL-27R signaling inactivation will result in heightened HCC development.

Here using two different *in vivo* models of HCC, one carcinogen-induced and injury-promoted and another NASH-driven, we surprisingly found that genetic inactivation of IL-27R suppresses HCC. Furthermore, through TCGA database analysis we found that elevated mRNA expression of IL-27R positively correlates with poor survival since the initial treatment alone and with more advanced stages of HCC development.

The emerging mechanism of immune-regulatory and pro-tumorigenic mechanism relies on the ability of IL-27R signaling to inhibit anti-cancer immune responses, particularly controlling NK cell accumulation and activation. Indeed, the experiment where NK cells were depleted during HCC development demonstrated that the effect of IL-27R signaling on HCC chiefly depends on NK cells. Liver microenvironment is uniquely enriched for NK cells, where they also exert immune surveillance (16). NK cells are potent killers of senescent, infected or cancerous cells and participate in regulation of immune responses via the production of inflammatory cytokines in the normal liver (16). During HCC development the number and activation of liver NK cells gradually reduces due to the chronic exposure to yet unidentified, presumably tumor-derived stimuli and reduction in expression of NK cell attracting chemokines (18). As obesity-induced non-alcohol steatohepatitis (NASH) is one of the major drivers of HCC, it is important to note that NK cell number and functions are also reduced in fatty liver disease (69).

We found that genetic ablation of IL-27R promoted the accumulation of NK cells in the tumor tissues, which was associated with NK-specific upregulation of CXCR6 in the tumor. Moreover, intra-tumoral NK cells exhibited heightened expression of cytotoxic molecules, implicating IL-27R signaling as a repressor of the NK-cell cytotoxic program. The activation of NK cells against target cells is dependent on the balance between activating and inhibitory receptors (43). We observed that NK cells isolated from tumors in the absence of competent IL-27R signaling exhibit upregulation of the NKG2D and the NKG2A activating receptors, which can “educate” hepatic NK cells, and downregulation of Ly-49C inhibitory receptor, which is more associated with education of peripheral NK cells. IL-27R signaling has been previously implicated in the regulation of NK cell function (70,71). For example, IL-27 together with IL-12 and IL-2 has been shown to promote NK cell activity (70). However, our data demonstrate that *in vitro* stimulation of NK cells with rIL-27 downregulates gene expression of cytotoxic molecules such as granule components (*Gzmb, Prf1*), FasL (*Faslg*) and IFN-γ (*Ifng*), as well as NKG2D (*Klrk1*) activating receptor. This together with *in vivo* observations of heightened cytotoxicity of NK cells in *Il27ra*^−/−^ HCC tumors suggest a direct suppressive function of IL-27R on NK cells.

Expression of so called ‘stress ligands’ on the surface of stressed or cancerous target cells as well as lack of MHC I expression is a second signal required to enable NK cell cytotoxicity (51,53). Our data revealed elevated expression of NK cell-stimulating stress ligands (*Raet1a*, *H60b*) and reduction in the *Tap1* MHC I-processing molecule and surface MHC I expression in HCC tumors of *Il27ra*^−/−^ mice. Moreover, the direct stimulation of HCC cells with rIL-27 suppressed *Raet1a* and *H60b* expression, implicating IL-27R signaling in the previously unknown control of HCC biology. Apart from direct regulation of NK cells, IL-27 could also be implicated in the regulation of other cell types, which in turn could mediate NK cell accumulation and activation. For instance, other immune cell subsets in the liver microenvironment could regulate NK cell function via cell contact interactions (PD-1, MHC I, 2B4-CD48) or via production of chemokines and cytokines affecting NK cell recruitment (CXCL9, CXCL10), their differentiation and/or maturation (IL-15, IL-12, IL-18) (72). In fact, we found that acute DEN administration triggered heightened mRNA expression of *Cxcl9* and *Cxcl10* chemokines and increased expression of the NK cell effector molecules perforin, granzyme B, TRAIL, and IFN-γ in the livers of IL-27R deficient mice 48 hours after DEN treatment. These observations are consistent with the scenario that IL-27R signaling represses the NK-mediated arm of cancer immunosurveillance, thereby enhancing HCC tumor initiation and progression.

With global obesity and type II diabetes epidemics, NASH-driven HCC is set to surpass all other forms of HCC in incidence (8,9). The *MUP-uPA* and Western Diet HCC models faithfully resembles human HCC driven by fatty liver disease and liver fibrosis, respectively. We found that IL-27R deficiency not only suppresses tumor development in this model, but also strongly reduces the underlying fibrosis. Similar to observations in the DEN model, *MUP-uPA*^+^*Il27ra*^−/−^ mice also showed enhanced NK cell activation, implying existence of a common mechanism regulated by IL-27R signaling. Interestingly, stellate cells and fibroblasts constituting fibrotic mass in NASH and HCC are often senescent (73) and NK cells have can be senolytic (74).

While several models of liver cancer are available, each of them has their advances and limitations (59). Our work, however, demonstrates that ablation of IL-27R limits liver cancer development in two different models of HCC: carcinogen-induced and injury-promoted HCC (DEN) and NASH-driven HCC (*MUP-uPA* +WD). This, along with human data on poor survival and advanced tumor stages in *Il27ra^hi^* HCC patients, implies that the IL-27 pathway plays a tumor-promoting role in HCC that is generalizable across different models and types of HCC with different drivers. An essential component of this mechanism is mediated through IL-27 dependent suppression of intra-tumoral NK cell function and increased innate resistance to tumors.

Taken together, our data uncover the important role of IL-27R-mediated regulation of NK cells in liver cancer development, where it directly and indirectly controls NK cell accumulation and activation, suggesting that inhibition of IL-27 or its receptor signaling could represent a novel preventive or therapeutic target in HCC.

## Methods

### Mice

*Il27ra*^−/−^ (JAX#018078) and C57BL6/J (WT) (JAX#000664) mice were purchased from Jackson laboratory and crossed to obtain *Il27ra^+/−^* and *Il27ra*^−/−^ mice. *Ncr1^gfp/gfp^* were obtained from Dr. Wayne Yokoyama (Washington University, St. Louis) and used to generate *Il27ra*^+/−^*Ncr1*^+/*gfp*^, and *Il27ra*^−/−^*Ncr1*^+/*gfp*^. For NASH model *Il27ra*^−/−^ mice were crossed to *MUP-uPA*^+^ (75) to obtain *MUP-uPA*^+^*Il27ra*^+/−^ and *MUP-uPA*^+^*Il27ra*^−/−^ mice. All mice were on C57BL/6 background. The genotyping was performed by standard PCR protocols. Mice were housed and bred under specific pathogen-free condition in an AAALAC-approved barrier facility at Fox Chase Cancer Center (FCCC). Littermate and cagemate controls were used for all experiments. All animal experiments were approved by the Institutional Animal Care and Use Committee (IACUC) at FCCC and performed in compliance with all relevant ethical regulations for animal research.

#### DEN model

To induce HCC development 25 mg/kg of diethylnitrosamine (DEN) (Sigma-Aldrich, N0258) was administered intraperitoneally into male mice at the age of 15 days as previously described (40). Mice were maintained on autoclaved water and a regular chow diet. HCC development was analyzed at 10 months of age.

For acute response mice were collected 48h after DEN (100mg/kg) administration.

#### NASH model

*MUP-uPA*^+^*Il27ra*^+/−^ and *MUP-uPA*^+^ *Il27ra*^−/−^ female and male mice were fed Western diet (Teklad, TD.88137) for 8 months beginning at 8 weeks after birth. HCC development was analyzed at 10 months of age.

#### Antibody treatment

200 μg/mouse of anti-NK1.1 (PK136, FCCC cell culture facility) or SV40 IgG2a (PAB419, FCCC cell culture facility) isotype control antibodies were injected i.p. weekly for 5.5 months starting from 4.5 months of age.

### Histology and Immunohistochemistry

For histological analysis, a liver lobe (with tumors) was isolated and fixed in 10% buffered formalin (Fisher HealthCare, 23-245685) for 24 h. Five μm thick sections were prepared and stained with hematoxylin (Sigma-Aldrich, HHS32) and eosin Y (Thermo Scientific, 6766007). All images were acquired with Nikon Eclipse 80i microscope.

For immunohistochemistry staining 5μm thick sections of livers containing tumors were deparaffinized by taking them through 4 changes in xylenes, then washed by 4 changes in 100% ethanol followed by re-hydration in tap water. Antigen retrieval was performed in 1X Citrate buffer (Electron Microscopy Sciences, 64142-08) at 95°C for 1 h followed by 1 h cooling down to RT. Then slides were rinsed in tap water for 3 min and dehydrated in 100% ethanol for 1 min, followed by blocking in 3% H2O2 in PBS for 10 min. Slides were blocked with 5% goat serum in 1% BSA-PBS for 20 min, then they were incubated with primary antibody for anti-Ki67 (1:100; BioLegend, 151202), anti-pERK1/2 (1:400; Cell Signaling, 4370) or anti-IL-27Rα (34N4G11; Novus Biologicals, NBP2-19015) O/N at 4°C. After washing with 1% BSA-PBS slides were incubated with secondary goat anti-rat and goat anti-rabbit biotinylated antibodies for 30 min at RT followed by 30 min of incubation with streptavidin–HRP (1:500; BD Pharmingen, 554066). For developing DAB substrate (Invitrogen) was applied for 3 min followed by washing in water and counterstaining with Hematoxylin solution (Sigma-Aldrich, HHS32). Excess of Hematoxylin was removed by immersing slides in 0.25% ammonia water followed by rinsing in water. Slides were mounted with coverslips using Permount mounting medium solution (Fisher Chemical, SP15). All images were acquired with Nikon Eclipse 80i microscope. Microsoft PowerPoint was used for one step brightness adjustment for all images in parallel. Quantification was done using ImageJ (version 1.51).

### Van Gieson staining

For collagen staining deparaffinized and re-hydrated slides were stained for 5 min in Van Gieson solution (EMS, 26374-06) followed by dehydration by 2 changes in 100% ethanol and cleared by 2 changes of xylenes. All images were acquired with Nikon Eclipse 80i microscope. Microsoft PowerPoint was used for one step brightness adjustment for all images in parallel. Quantification was done using ImageJ (version 1.51).

### Flow cytometry

Mice were sacrificed by CO2 inhalation and livers were perfused with HBSS containing 2% of Heparin (20 USP units/mL) to remove traces of blood. Livers were isolated, non-tumor and tumor tissues were dissected separately and incubated with a cocktail of digestion enzymes containing collagenase I (450 U/mL) (Sigma-Aldrich, C0130) and DNase I (120 U/mL) (Sigma-Aldrich, D4263) in HBSS (with Ca^2+^/Mg^2+^) for 40 min at 37°C with gentle shaking at 150 rpm. After incubation, cell suspension was filtered through a 70 μm cell strainer. Immune cells were enriched by density-gradient centrifugation over Percoll (GE Healthcare, 17-0891-01) at 1000xg for 25 min without brake (40% Percoll in RPMI-1640 and 80% Percoll in PBS). Leukocyte ring on a border of gradient and parenchymal cells on top were collected, washed and stained. The following antibodies were used: CD45-PercP (30F-11; BioLegend, 103130), CD11b-Pacific Blue (M1/70; BioLegend, 101224), NK1.1-FITC (PK136; BioLegend, 108706), NK1.1-PE (PK136; BioLegend, 108708), TCRβ-Alexa Fluor 700 (H57-597; BioLegend, 109224), CD4-APC/Cy7 (GK1.5; BioLegend, 100414), CD8α-APC (53-6.7; BioLegend, 100712), Ly6G-APC/Cy7 (1A8; BioLegend, 127624), Ly6C-PE/Cy7 (HK1.4; BioLegend, 128018), F4/80-APC (BM8; BioLegend, 123116), Granzyme B-Pacific Blue (GB11; BioLegend, 515408), CD27-PE/Cy7 (LG.3A10; BioLegend, 124216), Ly-49C-APC (4L03311), Ly-49I-PE (YLI-90; eBioscience, 1943023), NKG2AB6-APC (16A11; BioLegend, 142808), NKG2D-PE (CX5; BioLegend, 130208), CD49a-PE/Cy7 (HMα1; BioLegend, 142608), CD49b-APC/Cy7 (DX5; BioLegend,108920), CD11b-biotin (M1/70; BioLegend, 101204), CD31-PE (390; BioLegend, 102408), TER-119-biotin (TER-119; BioLegend, 1162204), H-2Kb-PE/Cy7 (AF6-88.5; BioLegend, 116519), H-2K^b^/H-2D^b^-APC/Fire 750 (28-8-6; BioLegend, 114617), Streptavidin-APC/Cy7 (BioLegend, 405208)

Spleens were isolated, mashed and filtered through 70 μm cell strainers. Peripheral blood was collected by cardiac puncture and erythrocytes were lysed by red blood cell (RBC) lysis buffer (15 mM NH_4_Cl, 0.1 mM NaHCO_3_, 0.1 mM Na_2_-EDTA) for 5 min at RT. Peripheral blood and splenocytes were stained with CD45-PercP (30F-11; BioLegend, 103130), NK1.1-FITC (PK136; BioLegend, 108706), NK1.1-PE (PK136; BioLegend, 108708), TCRβ-Alexa Fluor 700 (H57-597; BioLegend, 109224), Ly-49C-APC;, Ly-49I-PE (YLI-90; eBioscience, 1943023), NKG2AB6-APC (16A11; BioLegend, 142808), NKG2D-PE (CX5; BioLegend, 130208), CD49a-PE/Cy7 (HMα1; BioLegend, 142608) and CD49b-APC/Cy7 (DX5; BioLegend,108920). All antibodies were used at 1:50 dilution and LIVE/DEAD Fixable Yellow Dead Cell stain (Invitrogen, L34959) at 1:200.

### Gene expression analysis

Non-tumor and tumor tissues were homogenized in TRIzol reagent (Invitrogen, 15596018) with 2.8 mm ceramic beads (OMNI International, 19-646-3) using Bead Ruptor (OMNI International). Total RNA was extracted using Aurum Total RNA Fatty and Fibrous Tissue Kit (Bio-Rad, 7326870) according to manufacturer’s protocol. Sorted or treated cells were lysed in RLT Plus buffer (QIAGEN, 157030074) and total RNA was isolated using RNeasy Plus Mini Kit (QIAGEN, 74136) according to manufacturer’s protocol. Complementary DNA was synthesized using iScript Reverse Transcription Supermix (Bio-Rad, 1708841) with random primers according to manufacturer’s protocol. Q-RT-PCR was performed with Bio-Rad CFX 96 Connect Real-Time PCR Detection System using iTaq Universal SYBR Green Supermix (Bio-Rad, 1725124). The following primers were used: *Rpl32*, *Lcn2*, *Ccnd1*, *Cxcr6*, *Gzmb*, *Prf1*, *Tnfsf10*, *Faslg*, *Ifng*, *Klrk1*, *Raet1a*, *H60b*, *Tap1*, *Cxcl9*, *Cxcl10*.

### Clonogenic assay

1000 DEN-derived HCC cells were plated on 0.1%-swine gelatin precoated 6-well plate in ACL-4 containing 20% FBS in triplicate per condition. 24 h later medium was changed to ACL-4 containing 5% FBS with or without rIL-27 (200 ng/ml). On day 4 medium was refreshed and cells were left to grow for additional 3 days. On day 7 after beginning of the treatment the medium was aspirated, cells were washed with HBSS (Ca^2+^/Mg^2+^ free) and fixed 10% acetic and 10% methanol fixing solution for 15 min at RT. When fixing solution was aspirated, plates were left to dry followed by adding 0.4% crystal violet staining solution for 20 min at RT. Rinsed with tap water wells were scanned with EPSON Perfection V600 Photo scanner.

### Immunomagnetic purification of NK cells and treatment with rIL-27 *in vitro*

NK cells were purified from spleen of wild type male mice by negative selection using magnetic beads and a cocktail of biotinylated antibodies: CD3e (145-2C11; BioLegend, 100304), CD4 (GK1.5; BioLegend,100404), CD8α (53-6.7; BioLegend, 100704), CD11b (M1/70; BioLegend, 101204), CD11c (N418; BioLegend, 117304), Gr-1 (RB6-8C5; BioLegend, 108404), TER-119 (TER-119; BioLegend, 1162204), B220 (RA3-6B2; BioLegend, 103204) and CD19 (6D5; BioLegend, 115504). Purified NK cells were plated at a density of 250,000 cells/mL in 2 mL of complete RPMI-1640 without FBS containing 25 ng/mL of rIL-27. 12 h later cells were lysed and used for gene expression analysis.

### HCC stimulation *in vitro*

For gene expression analysis 500,000 DEN-derived HCC cells were plated on a 0.1%-swine gelatin precoated 6-well plate in ACL-4 containing 20% FBS in triplicate per condition. 24 h later medium was changed to complete DMEM without FBS with or without rIL-27 (200 ng/ml) for 3 h at 37°C in a 5% CO_2_ cell culture incubator. For Western blot analysis cell were treated as described above and collected at 0, 15 min, 30 min.

### Partial hepatectomy

2/3 partial hepatectomy was performed as previously described (76). Briefly, mice were anesthetized by isoflurane. Skin and abdominal wall were cut open; left and right lobe were ligated sequentially with silk suture and excised. Then abdominal wall was sutured and skin was clipped. Liver regeneration was analyzed in 8 days.

### Western Blot

*In vitro* treated HCC were washed with HBSS (Ca^2+^/Mg^2+^ free) and lysed in RIPA buffer supplemented with phosphatase and protease inhibitors (Sigma-Aldrich, PPC1010) (200 μL per 10^6^ cells) followed by centrifugation at 15,000xg for 10 min at 4°C to pellet not lysed cell debris. The supernatant was collected for the analysis. Protein concentration was determined by BCA Protein Assay Kit (Sigma-Aldrich, 1001491004) according to manufacturer’s protocol. 40 μg of cell lysates was separated by 4-20% Tris-glycine MINI-PROTEAN TGX gels (Bio-Rad, 456-1094) and transferred to PVD membranes using Trans-Blot Turbo Transfer Pack (Bio-Rad, 1704156). Each membrane was washed with TBST (10 mM Tris-HCl, 150 mM NaCl, 0.1% Tween-20; pH 7.6) and blocked with 5% skimmed milk in TBST for 1 h followed by O/N incubation at 4 °C with appropriate primary antibody: pSTAT3 (D3A7; Cell Signaling Technology, 9145S) and STAT3 (124H6; Cell Signaling Technology, 9139S). Loading was evaluated by staining with β-actin-horseradish peroxidase (HRP) (AC-15; Abcam, ab49900) antibody (1:50000) for 1 h at RT. Each membrane was washed and primary antibodies (except β-actin-HRP) were detected with a 1:5000 dilution of HRP-conjugated rabbit anti-mouse IgG (Cell Signaling Technology, 7076S) or mouse anti-rabbit IgG (Cell Signaling Technology, 7074S). The reactive bands were developed using ECL Prime western blotting detection reagent (GE Healthcare, RPN2232) and were visualized with an autoradiography film (LabScientific, XAR ALF 2025).

### NK cytotoxicity *in vivo*

NK-cell cytotoxicity was measured as previously described (77). Briefly, RMA and RMA-S T-cell lymphoma cells were labelled with Orange CMRA (Invitrogen, C34551) or CPD eFluor 650 (eBioscience, 65-0840-90) dyes, respectively. 2×10^5^ cells of each cell line were mixed in 1:1 ratio and injected i.p. to *Il27ra*^+/−^ or *Il27ra*^−/−^ mice. 48h later mice were sacrificed, peritoneal lavage was collected and analyzed by FACS.

### NanoString

50 ng of tumor RNA was used for NanoString in order to analyze immune profile of the tumor microenvironment measuring the expression of 770 genes according to manufacturer’s protocol. The hybridization between target mRNA and reporter-capture probe pairs was performed at 65°C for 20 h using Applied Biosystems Veriti Thermal Cycler. All processing was carried out on a fully automated nCounter Prep Station. Excess of probes was removed and probe-target complexes were aligned and immobilized in the nCounter cartridge followed by the image acquisition and data processing by nCounter Digital Analyzer. The expression level of a gene was measured by counting the number of times the specific barcode for that gene was detected, and the barcode counts were then tabulated in a comma-separated value (CSV) format. The raw digital count of expression was exported from nSolver v3.0 software. Statically significant differentially expressed genes between genotypes were analyzed by KEGG pathway analysis.

### Statistical Analysis

Student’s two-tailed *t*-test was used for comparison between two groups. Survival curve data were analyzed using the long-rank (Mantel-Cox) test. Chi-square test was used to compare stages of human HCC. Tukey’s test was used for multiple comparisons. Data were analyzed using the GraphPad Prism Software (Version 7.0). Data are presented as mean ± SEM; *p < 0.05, **p < 0.01, ***p < 0.001, ****p<0.0001. A p-value < 0.05 was considered statistically significant.

## Authorship Contributions

T.A., S.I.G and E.K.K. designed the study, planned the experiments, analyzed the data. T.A., I.O.P., A.M.M., E.K.T, and A.F.R. performed the experiments. T.A., K.S.C., S.I.G., and E.K.K. wrote the manuscript.

## Acknowledgments

We acknowledge the help of FCCC facilities as well as Dr. Ofer Mandelboim (Hebrew University, Jerusalem) for access to Ncr1^gfp/gfp^ mice and Dr. Wayne Yokoyama (Washington University, St. Louis) for providing the mice after backcross into the C57BL/6 background and microsatellite mapping verification. This work was supported NIH/NCI Cancer Center Support Grant P30 CA006927 to Fox Chase Cancer Center; WW Smith Charitable Trust, NIH R21 CA202396, R01 HL133669 and R01 HL149946 grants to E.K.K, NIH R01 CA227629 and CA218133 to S.I.G

## Supplemental Figures

**Supplemental Figure 1.**
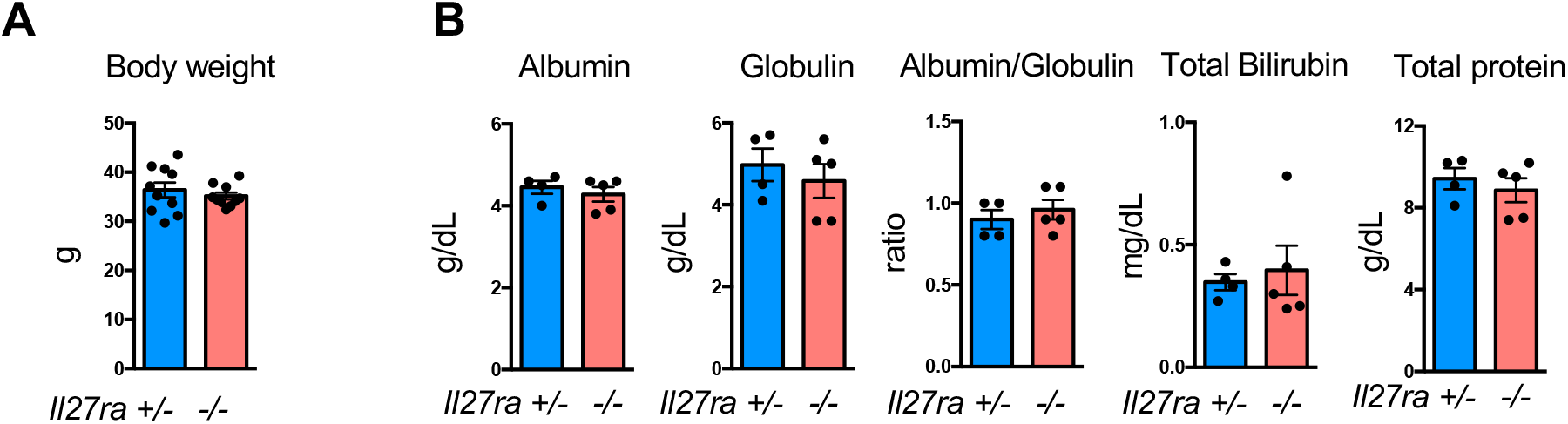
IL-27R deficiency does not affect body weight and serum proteins. **(A)** Body weight of DEN-injected tumor-bearing *Il27ra*^+/−^ (n=10) and *Il27ra*^−/−^ (n=10) male mice. **(B)** Serum analysis of DEN-injected tumor-bearing *Il27ra*^+/−^ (n=4) and *Il27ra*^−/−^ (n=5) male mice. Data are mean ± SEM from at least 3 independent experiments from 10-month-old mice.

**Supplemental Figure 2.**
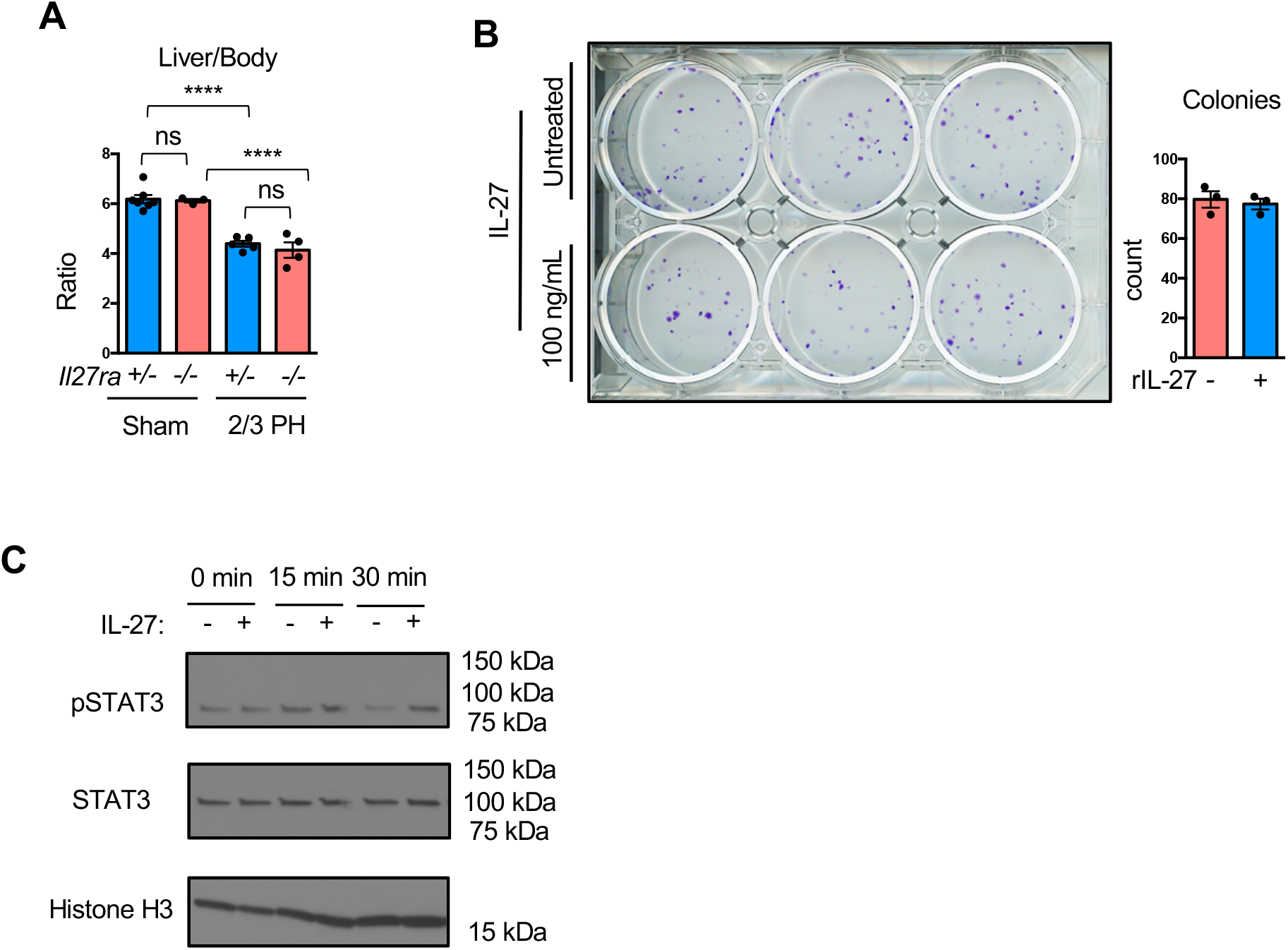
IL-27R deficiency does not alter liver regeneration and colony forming ability of HCC cells. **(A)** Liver to body weight ratio of *Il27ra*^+/−^ (n=7) and *Il27ra*^−/−^ (n=3) male mice sham surgery and *Il27ra*^+/−^ (n=5) and *Il27ra*^−/−^ (n=4) male mice subjected to 2/3 partial hepatectomy. **(B)** Representative image and quantification of the colonies grown from DEN-derived HCC cells treated *in vitro* with rIL-27. Data are mean ± SEM from at least 3 independent experiments. ****p<0.0001, Tukey’s multiple comparisons test. **(C)** pSTAT3 and STAT3 protein expression analysis in protein lysates of DEN-derived HCC cells treated with rIL-27 and collected at different time points (0, 15, 30 min).

**Supplemental Figure 3.**
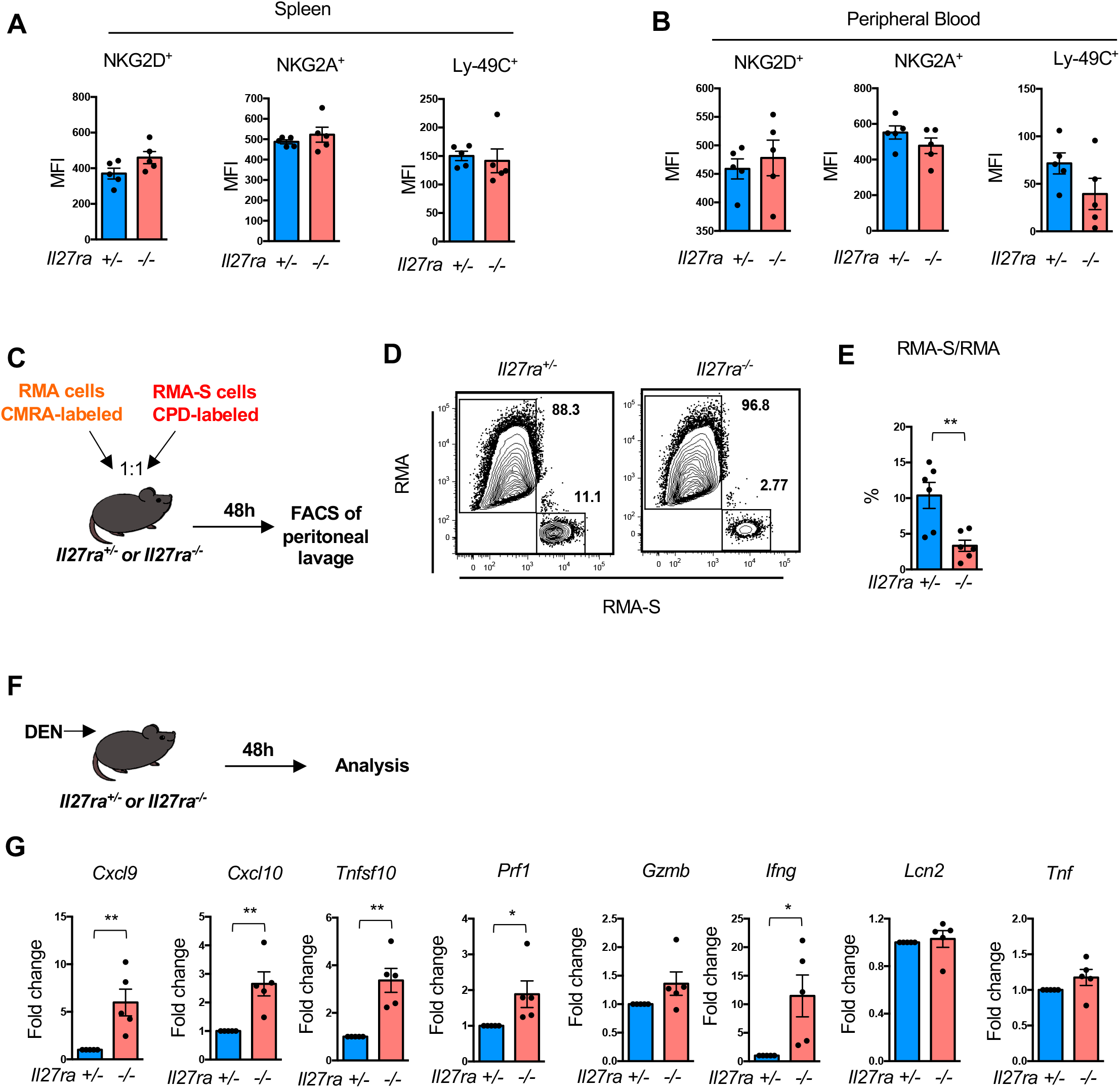
IL-27R signaling is implicated into regulation of NK cell activity. Single cell suspensions of spleen and peripheral blood from DEN-*Il27ra*^+/−^ (n=5) and *Il27ra*^−/−^ (n=5) 10-month-old mice were stained for Live/Dead, CD45, TCRβ, NK1.1, CD49a, CD49b, NKG2D, NKG2A and analyzed by FACS. Mean fluorescence intensity of NKG2D^+^CD49b^+^, NKG2A^+^CD49b^+^ and Ly-49C^+^CD49b^+^ NK cells in spleens (A) or peripheral blood (B). **(C)** Scheme of the experiment. Cell lines were dye-labelled, mixed in 1:1 ratio and i.p. injected into 8-week-old *Il27ra*^+/−^ and *Il27ra*^−/−^ mice, cell recovery was analyzed in 48 h. **(D)** Representative plots of RMA and RMA-S, sensitive and insensitive to NK cells killing, respectively. (E) Percentage of RMA /RMA-S cells ratio from peritoneal lavage of *Il27ra*^+/−^ (n=6) and *Il27ra*^−/−^ (n=6) mice. **(F)** Scheme of the experiment. 8-week-old *Il27ra*^+/−^ and *Il27ra*^−/−^ male mice were injected with 100 mg/kg of DEN, livers were collected for analysis in 48 h. **(G)** Relative gene expression of chemokines *Cxcl9*, *Cxcl10*, and cytotoxic molecules *Tnfsf10*, *Prf1*, *Gzmb* and *Ifng* in livers from *Il27ra*^+/−^ (n=5) and *Il27ra*^−/−^ (n=5) male mice. Gene expression was first normalized to *Rpl32* then to gene expression in liver of *Il27ra*^+/−^ mice. Data are mean ± SEM from at least 3 independent experiments. *p<0.05, **p<0.01, unpaired Student’s t-test (two-tailed).

**Supplemental Figure 4.**
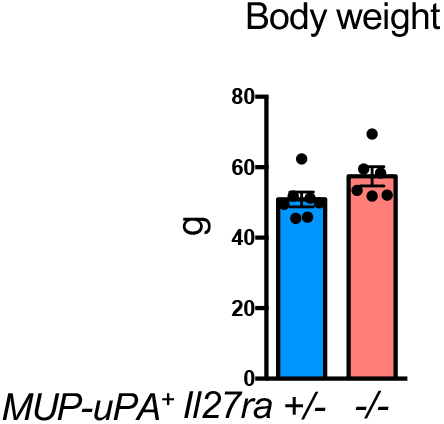
IL-27R deficiency does affect body weight of mice in NASH-dependent model of HCC. Body weight of WD-fed tumor-bearing mice *MUP-uPA*^+^*Il27ra*^+/−^ (n=7) and *MUP-uPA*^+^*Il27ra*^−/−^ (n=6) male mice. Data are mean ± SEM from at least 3 independent experiments.

**Supplemental Figure 5.**
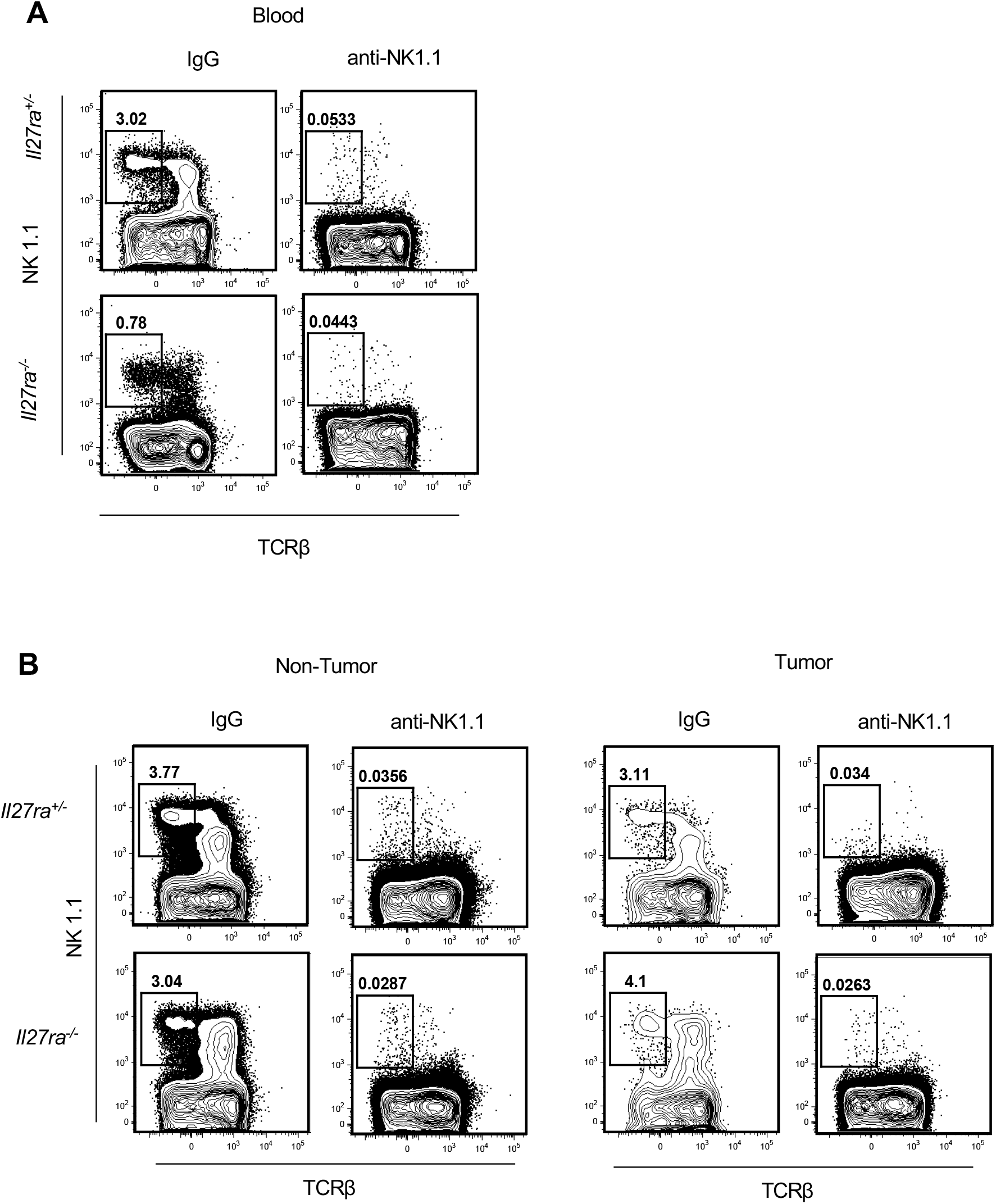
Efficiency of long-term NK-cell depletion. Representative FACS plots of NK cell depletion efficiency in blood (A) and non-tumor and tumor tissue (B) from *Il27ra*^+/−^ and *Il27ra*^−/−^ male mice which received anti-NK1.1 antibody or isotype control. Data are mean ± SEM from at least 3 independent experiments.

